# Brain-Body Interactions Influence the Transition from Mind Wandering to Awareness of Ongoing Thought

**DOI:** 10.1101/2024.09.03.610929

**Authors:** Kazushi Shinagawa, Yuto Tanaka, Yuri Terasawa, Satoshi Umeda

## Abstract

Our thoughts are inherently dynamic and often wander far from our current situation (mind wandering, MW). Although previous research revealed that the ascending arousal system shapes neural dynamics to mediate awareness of ongoing thoughts, the physiological states and afferent signals altered by this activation and its effects on awareness are unknown. In this study, we examined electroencephalography (EEG), electrocardiography (ECG), and respiration data before participants were aware of MW during a task in which they focused on external or internal stimuli. We showed that the transition from MW to awareness was characterized by decreased alpha and beta activity and increased heartbeat-evoked potential (HEP) amplitudes. In addition, the participants were more likely to be in the exhalation phase becoming aware, and in the inhalation phase at the time of MW reports. Moreover, changes in cardiac activity and HEP accompanied this pattern when participants were asked to focus on respiration. Based on these findings, we suggest that the release from the increased cognitive load with sustained MW and catching these changes as physiological alterations supporting awareness of MW; moreover, the modulation of the respiratory cycle by focusing on breathing enhances these changes.

## Introduction

The human brain dynamically generates conscious experiences, encompassing various aspects of our subjective experiences. Humans spend a large proportion of these experiences engaged in mind wandering (MW^1,2^), which is often defined as self-generated thoughts that are decoupled from the current environment or task^3^. To reduce the time spent on MW in undesirable situations (e.g., at work or in class), it is important to become aware of MW immediately after it occurs^4^. The inability to monitor and disengage from excessive or distracting self-generated thoughts is associated with impairments in well-being^5^. However, the transition from MW to the awareness of ongoing thoughts, a kind of change in conscious experience, is not yet fully understood.

In our daily lives, the brain generates cognitive experiences both on its own and through interactions with the body^6,7^. Previous research on the dynamics of conscious experience has focused on brain-body interactions^8–11^. Empirical evidence has revealed that the autonomic nervous system and respiratory phases modulate information processing throughout the brain^12,13^. In addition to effects at the system level, information from the body itself influences conscious experiences, including thought. The relationship between the variability and self-relevance of thought with the variability in cardiac activity and its central process has been experimentally demonstrated respectively^14,15^. The heartbeat-evoked responses in electroencephalography (EEG) data are more accurate for classifying unconsciousness and consciousness than the use of random EEG segments^16^. Considering the evidence that cardiac activity modulates perception and emotion^7,17,18^, afferent signals from cardiac activity may contribute to various aspects of conscious experiences. Additionally, respiration is closely related to the locus coeruleus (LC), which supports sustained attention^19^. Respiration affects the heart rate physiologically, and the central process of cardiac activity is modulated by respiration phases^20^. The relationships among neural activity, cardiac activity, respiration and consciousness highlight the potential effects of brain-body interactions on conscious experiences, including the awareness of ongoing thoughts at the system and content levels.

Although there are no direct findings, previous research on MW has suggested the role of physiological activity in forming awareness of MW. In a recent study, changes in neural activity during conscious experiences were tracked via the self-caught method^12^, in which participants reported whenever they noticed MW^21^. With this method, the changes in thoughts from MW to awareness are extracted and reported. The results revealed that the ascending arousal system is activated several seconds before MW is reported^12^. This system is assumed to dynamically shape the excitability and receptivity of neurons across the brain rather than content specificity^22^. These results suggest that sympathetic activity can regulate ongoing brain dynamics at the system level, supporting changes in conscious experiences^8^. However, as the sympathetic activity does not immediately lead to MW, the detailed processes by which sympathetic activity results in the awareness and subsequently reports of MW remain unknown. Furthermore, sympathetic activity leads to alterations in brain activity at the system level. While awareness of ongoing thought is also recognized as a change in the content of thought, content-level processes remain poorly understood. To address this gap and clarify the mechanisms involved in becoming aware of ongoing thought, we investigated the dynamics of thoughts, and the brain and body activities mediated by the sympathetic nervous system.

In the present study, we aimed to investigate the dynamics of brain activity and the physiological state that leads to awareness. We combined a simple reaction task with the self-caught method and recorded EEG, electrocardiogram (ECG), and respiration data (Fig. 1a-c). The reaction task included two conditions: the participants responded continuously to the presented sounds (sound-focused; SF) or their exhalations (breathing-focused; BF). The acquired data before and after MW reports were segmented into three states to investigate changes related to awareness (Fig. 1c). The MW state, in which the participants were immersed in their internal thoughts, was defined 5–9 seconds before the reports of MW on the basis of previous studies^23–25^. The aware state was defined as the duration during which brain activity and the physiological state transitioned from MW to awareness, which occurred 1 to 5 seconds before MW was reported. These definitions are based on the finding that increased sympathetic activity and external perceptual processing, which may reflect leaving from the MW, occurs 4 or 3-5 seconds before MW is reported^12,26^. To characterize the MW state, we defined the focus state as the duration during which the participants concentrated on external stimuli (SF) and respiration (BF) when the first sound stimulus was presented after resuming the task for 4 seconds. The time after the task was resumed was defined as the interval of concentration to the task on the basis of previous studies^23–25^. These time windows were also determined to ensure a sufficient time window to calculate each of the indicators.

**Fig. 1.**
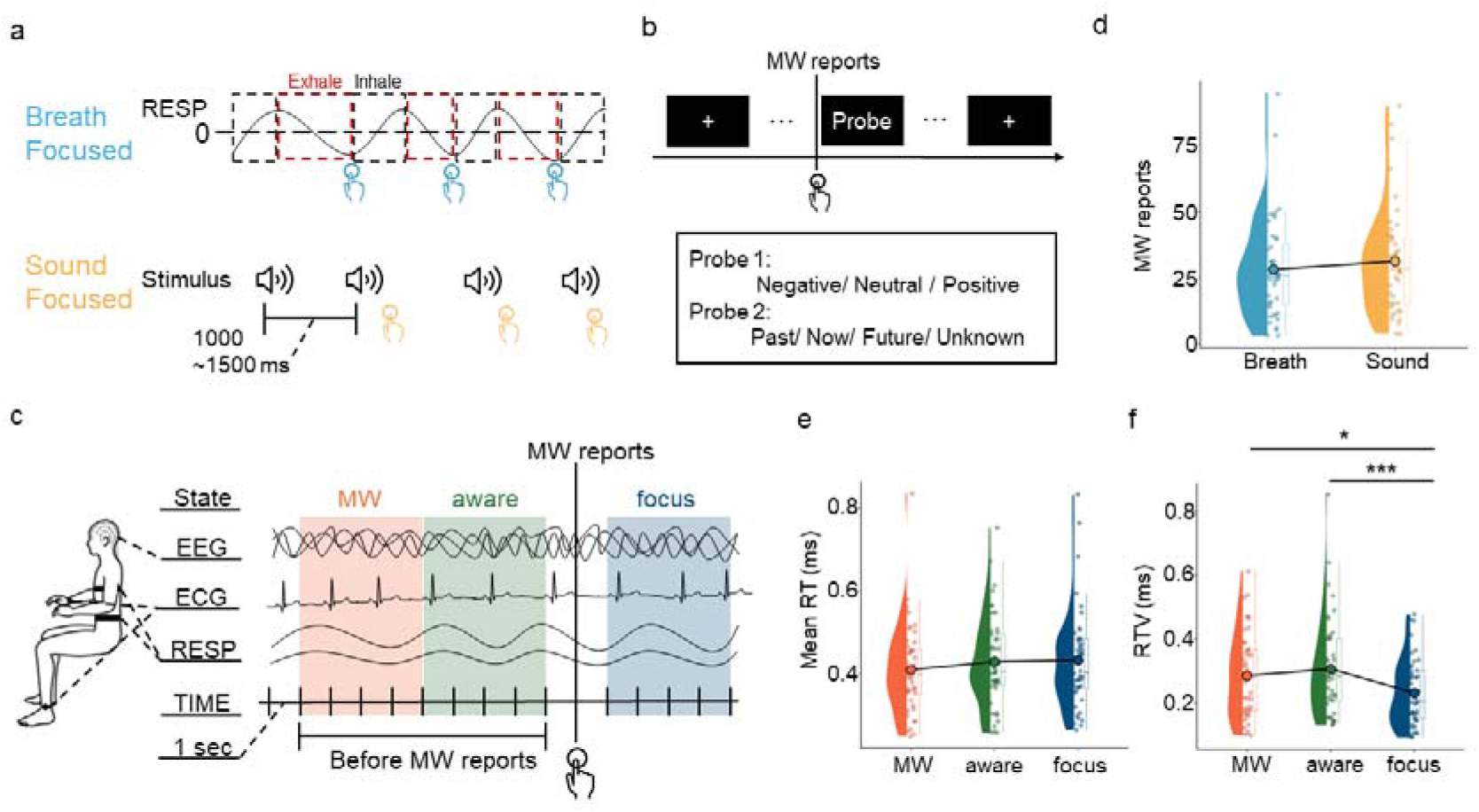
Experimental settings and behavioural results. **a, b. Overview of the experimental design:** In the breathing-focused condition, the participants responded at the end of exhalation, and in the sound-focused condition, they responded to each sound. During the response task, participants reported when they became aware of MW and answered questions about the content of their thoughts (emotional valence and time orientation) before resuming the task. Sound stimuli were consistently presented in both conditions. Each task was performed for four 12-minute sessions. **c. Measurement indices and definitions of attentional states:** We recorded EEG (65 channels), ECG (1 channel), and respiration (2 channels) data during the task. The interval from 1 to 5 seconds before MW reports defined the aware state, whereas the period from 5 to 9 seconds before MW was reported was defined as the MW state. The focus state was defined as the 4 seconds after the initial sound stimulus after the task was resumed. **d. MW frequency per task condition:** The horizontal axis shows the task condition, and the vertical axis shows the average number of MW reports. Each dot represents the data for one participant. **e, f. Reaction time and its variability among attentional states:** The horizontal axis shows the attentional state, and each dot represents the mean value for one participant. \**p* < .05, ** *p* < .01, *** *p* < .001.

To clarify the mechanism underlying spontaneous awareness of MW, we focused mainly on neural activity, the physiological state, and the central process of cardiac activity in the above thought states. Time-frequency analysis was performed based on the EEG data to examine the influence of overall neural activity on awareness of MW. In addition, we examined physiological states from cardiac activity and respiratory phases, which influence the conscious experience of thoughts. As indices of these central processes, we utilized heartbeat-evoked potentials (HEP), which are EEG components obtained by averaging at the R peak of the heartbeat^27^. The HEP is considered to reflect afferent signals from the heartbeat and processing in the brain, and many studies have utilized HEP to investigate how the process of cardiac activity influences cognitive functions^28^. A recent study revealed that heartbeat-induced pulsations of cerebral blood vessels can directly affect central neuronal activity by activating mechanosensitive channels, supporting the physiological validation of HEP^29^. By comparing these indices in attentional states, we investigate how brain–body dynamics lead to awareness.

## Results

### Subjective experiences and task performance

In the present study, we recruited 44 healthy university students in Tokyo with normal or corrected-to-normal vision and hearing (24 women; mean age = 21.21; *SD* = 1.96; *range* = 18-29). The average number of MW reports was 28.39 (*SD* = 19.22, *range* = 3-96) in the BF condition and 31.70 (*SD* = 20.51, *range* = 4-90) in the SF condition (Fig. 1d). First, task performance was analysed to confirm that the MW reports obtained in this study showed the same trends as those reported in previous studies. We compared the reaction time (RT) and the response time variability (RTV), which were calculated by dividing the RT variance within each interval the mean value in the focus and MW states. Although the RT did not significantly differ between the focus and MW states, a significant difference was found for the RTV (*F*(2, 86) = 8.365, *p* < .001, η*^2^* = .16). Subsequent analyses revealed less variability during the focus state than during the MW and aware states (MW: *t*(86) = 2.837, *p* = .0155, *d*= .62; aware: *t*(86) = 3.970, *p* < .001, *d* = .86; Fig. 1e and f). Consistent with previous research^30,31^, our study revealed significant variability in the responses in the MW state.

### Changes in whole-brain activity in the transition from MW towards awareness

Next, we clarified how the brain dynamics were characterized during the transition from the MW to the Aware state. Using a cluster-based permutation test (see “Methods”), we compared the spectral power across various frequencies at each electrode between the MW and focus states (Fig. 2a). Assuming a consistent attentional state within this interval, we averaged the data over time, yielding values for each channel by frequency. In Fig. 2a, warmer colours denote higher intensity in certain frequency bands during MW, whereas cooler colours indicate greater intensity in the focus state. The dotted lines indicate areas identified as significant in the cluster analysis (*p* <. 025). Our analysis revealed a uniform trend across both task conditions: theta (below 7 Hz), beta (13 to 30 Hz), and gamma (above 30 Hz) wave activities were higher in the focus state than in the MW state (BF: *t_cluster_*(43) = −3516, *p* <.001; SF: *t_cluster_* (43) = −3769, *p* <.001). In addition, our analysis revealed increased alpha wave (7 to 13 Hz) activity during MW, although this trend was not significant in the cluster analysis. The frequency maps of the MW states, when compared to the Focus state, are similar to those of previous studies^32^, supporting the validity of the MW states identified in the present study.

**Fig. 2.**
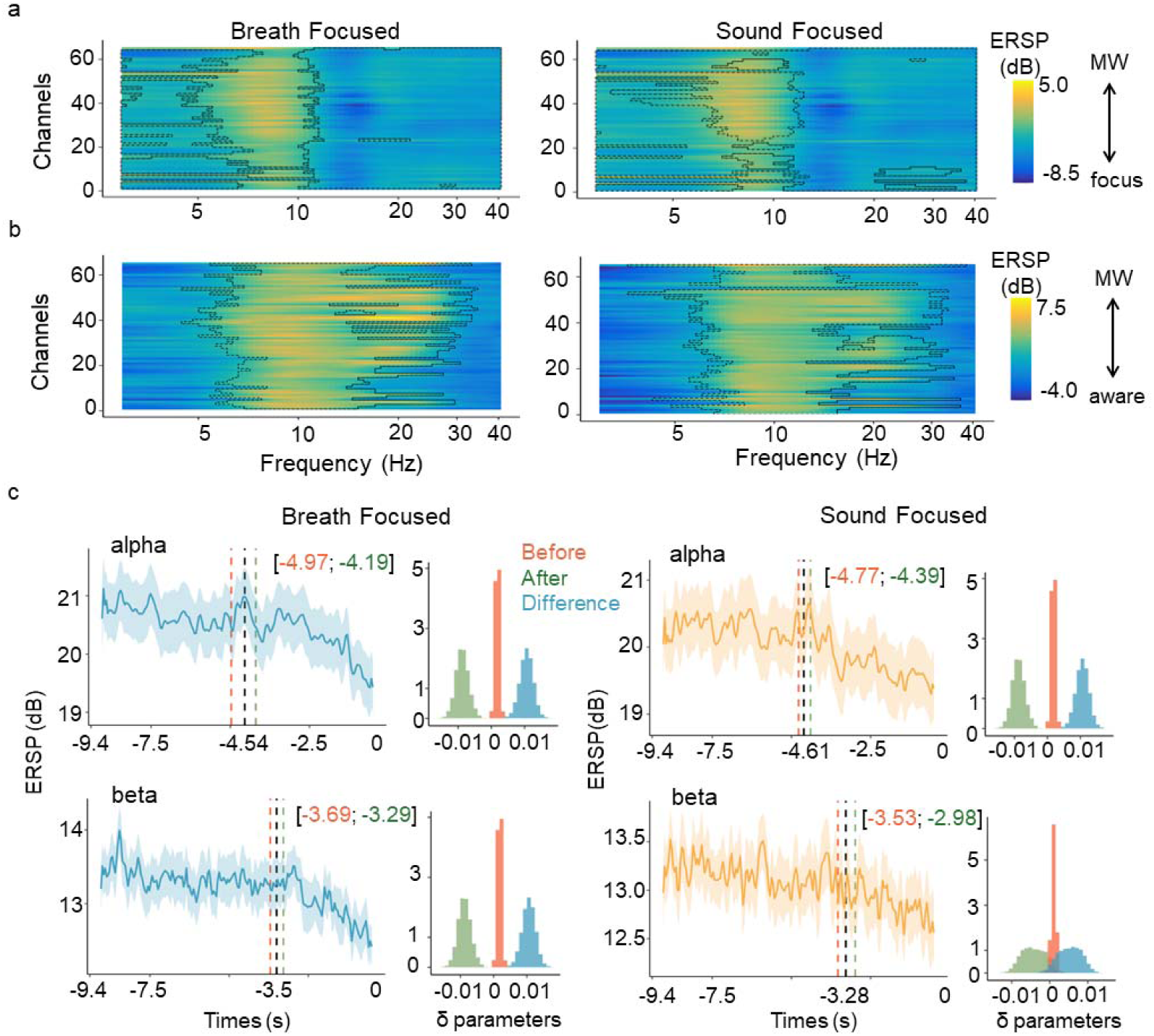
Changes in time-frequency components related to awareness. **a. Difference in frequency intensity between the MW and focus states:** We show the statistical value of the difference between the frequency of each electrode in the MW and focus states. The vertical axis shows the electrode number, and the horizontal axis shows the frequency band. Warmer colours indicate greater intensity in certain frequency bands during MW, whereas cooler colours indicate greater intensity in the focus state. The dotted lines mark areas identified as significant in the cluster analysis. **b. Difference in the frequency intensity between the MW and aware states:** Same as in **a**, except for the state being compared. Warmer and cooler colours indicate increased intensity in the MW and aware states, respectively. **c. Time series of the spectrum power before MW reports:** The horizontal axis represents the time in seconds from when MW is reported, and the vertical axis represents the spectral power values. The bold line represents the mean value, and the range represents the standard error. The vertical dotted line represents the 95% credible interval (CI) of the estimated change point. The interval before the estimated change point is until the red line, and the interval after the change point is on the right side of the green line. The trends before and after the change point are plotted on the right side of the time series. The red and green histograms show the distributions of the trend parameters before and after the change point, respectively. The difference parameter indicates the value obtained by subtracting the trend after the change point from the value before the change point in the blue histogram.

Additionally, we analysed the frequency differences between the MW and aware states, and the results are presented in Fig. 2b. This figure shows the differences in intensity for each frequency band and channel, with warmer and cooler colours indicating increased intensity in the MW and aware states, respectively (Fig. 2b). The results demonstrated that alpha and beta band activities were lower in the aware state than in the MW state (BF: *t_cluster_* (43) = 2162, *p* = .003, SF: *t_cluster_* (43) = 1935, *p* = .006).

Based on the cluster-based permutation test, we identified when alterations in alpha and beta waves began and clarified the sequential relationship between these changes using the time series analysis with change point detection (see “Methods”). The averaged time series of each frequency band at the Cz electrode, the electrode located in the centre, was used to estimate the model parameters (Fig. 2c). For the alpha wave, the analysis highlighted a 95% credible interval (CI) for the change points, which indicates the starting point of a decrease trend, ranging from 4.97 to 4.19 seconds before MW reports in BF condition. This result suggests a 95% probability that a change in the time series occurred within this interval. The estimated mean change point was 4.67 seconds before MW was reported (SF: mean = 4.61, 95% CI = [4.39; 4.77]). We plotted the trend parameters before and after the change points on the right side of the time series in Fig. 2c. A comparison of the mean values of the trends before and after the change point revealed that the lower limit of the 95% CI exceeded 0, indicating that alpha wave activity turned to a decreasing trend after the change point (BF: mean = 0.00984, 95% CI = [0.00592; 0.0132]; SF: mean = 0.152, 95% CI = [0.0104;0.0192]). Similarly, the change point for the beta wave activity was estimated to occur 3.69 to 3.29 seconds before MW was reported with a 95% probability (BF: mean = 3.51 seconds before the report; SF: mean = 3.28, 95% CI = [2.98; 3.53]). A comparison of the mean values of the trends before and after the change point revealed that the lower limit of the 95% CI exceeded 0, indicating that beta wave activity decreased after the change point (BF: mean = 0.0105, 95% CI = [0.00717; 0.0137]; SF: mean = 0.005, 95% CI = [0.0002;0.0101]). The change point analysis suggests that the decrease in beta wave activity typically occurs after the decrease in alpha wave activity.

### Changes in physiological states and their central process in the transition from MW toward awareness

#### Interbeat interval as an index of physiological states

To examine the effect of sympathetic nervous system activity on the physiological state, we analysed the effect of the attentional state on the inter-heartbeat interval (RR interval; Fig. 3a). While task conditions did not significantly influence the RR interval, a significant main effect of the attentional state was found (*F*(2, 212.02) = 29.915, *p* < .001, η*^2^*=.22). The subsequent analysis revealed that the heart rate was faster in the focus state than in the MW and aware states in each condition (MW: *t*(212) = 6.762, *p* < .001, *d* = .93; aware: *t*(212) = 6.634, *p* < .001, *d* = .91; Fig. 3c). Importantly, no significant differences in heart rate were found between the MW and aware states. A previous study noted MW was associated with low arousal in response tasks to external stimuli^33^.

**Fig. 3.**
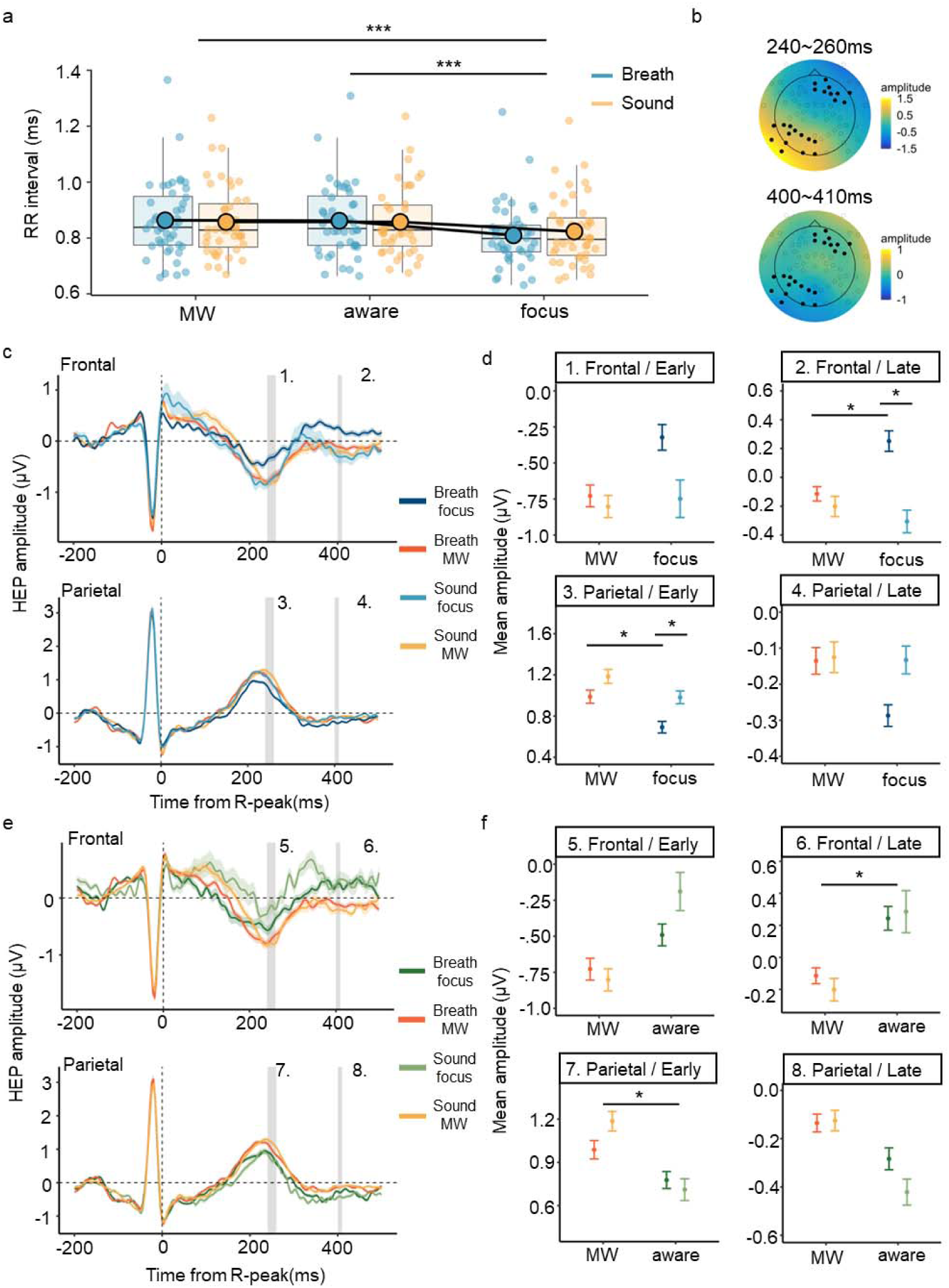
Cardiac activity and its central process in each task condition and attentional state. **a. Cardiac activity in each task condition and attentional state:** The horizontal axis shows the attentional state, the vertical axis shows the RR interval, and the colour indicates the task condition. Each dot indicates a participant. **b. Topographical map across time intervals:** In this map, colour gradients are used to depict an amplitude map—warmer colours represent higher potentials, whereas cooler colours indicate more negative potentials. We plotted the data collected at all the electrodes in a circle, and the electrodes used for heartbeat-evoked potential (HEP) analysis are indicated with black dots. **c. HEP waveforms in the MW and aware states:** The average HEP waveforms from the frontal and parietal electrodes are shown in the upper and lower panel, respectively. The vertical axis shows the amplitude, and the horizontal axis shows the time from the R peak. The bold line shows the mean value, and the range shows the standard errors. The mean HEP amplitude for each participant, focusing on early and late components, was computed by averaging across the periods highlighted in grey. **d. Mean HEP amplitude at the frontal and parietal electrodes:** We present the results from the frontal and parietal electrodes in the upper and lower panels. The early component (240~260 ms) and late component (400~410 ms) results are displayed on the left and right sides, respectively. The vertical axis shows the mean amplitudes, the horizontal axis shows the attentional state, and the colour indicates the task condition. The error bars indicate the standard errors. **e, f. HEP amplitudes in the aware and MW states:** Same as in **c, d**, except for the states being compared. * *p* < .05, ** *p* < .01, *** *p* < .001.

#### Heartbeat-evoked potentials as indices of the central process of cardiac activity

We analysed the mean HEP amplitude across task conditions and attentional states to explore how the central process of cardiac activity changes in the transition from MW to awareness (Fig. 3b-f). The mean HEP amplitudes were calculated from frontal and parietal electrodes for the early (240~260 ms) and late components (400~410 ms). First, we compared the mean HEP amplitudes in the MW and focus states to clarify how the signals from the body changed during MW. The analysis using the frontal electrodes revealed a significant main effect of the task conditions and a trend of interaction in the late components (*F*(1, 111) = 6.278, *p* = .013, η*^2^* = .05; *F*(1, 111) = 3.35, *p* = .069, η*^2^* = .03). Subsequent analysis indicated that HEP amplitudes were significantly greater during the Focus state than during the MW state in the BF condition (*t*(111) = 2.012, *p* = .046, *d* = .38). Moreover, a comparison between conditions in the focus state revealed higher HEP amplitudes in the BF condition than in the SF condition (*t*(111) = 3.063, *p* = .002, *d* = .50). A pattern similar to the late component at the frontal electrode was observed for the early component at the parietal electrode, with significant effects noted for both the task condition and the attentional state (*F*(1, 111) = 5.598, *p* = .019, η*^2^* = .05; *F*(1, 111) = 3.350, *p* = .017, η*^2^* = .05; Fig. 3c and d). Similar to the frontal electrode analysis, the HEP amplitudes in the focus state were lower than those in the MW state in the BF condition (*t*(111) = −2.031, *p* = .044, *d* = −.39), and lower in the BF condition than in the SF condition (*t*(111) = −1.992, *p* = .048, *d* = −.38).

Next, we explored how the central process of cardiac activity during MW changes in the transition towards awareness (Fig. 3e and f). The analysis for the late component at the frontal electrode revealed a significant main effect of the attentional state (*F*(1, 111) = 8.068, *p* = .005, η*^2^* = .06), whereas the main effect of the task condition and the interaction effects were not significant. Similarly, for the early component at the parietal electrode, only a main effect of the attentional state was observed, with increased HEP amplitudes in the aware state compared with those in the MW state (*F*(1, 111) = 7.302, *p* = .008, η*^2^* = .06). We found no other significant effects. These results suggest that participants in the aware state showed increased the central process of cardiac activity, similar to that observed when they focused on breathing, compared to that in the MW state.

### Respiratory phases and their effects on cardiac activity and its central process in the transition from MW towards awareness

In our final analysis of the transition from MW towards awareness, the respiratory phases and their effects on cardiac activity and HEP before MW was reported were examined (Fig. 4). First, we revealed that the respiratory cycle was decreased in the BF condition, suggesting slower respiration than in the SF condition (*t*(43) = −11.105, *p* < .001, *d* = 3.39; Fig. 4a). Next, we examined whether there is the pattern of respiratory phases corresponds to the spontaneous awareness. For this purpose, we tested whether the rate of the inhalation phase every second before MW reports were more or less than the baseline, which was calculated as the mean value of 10000 samples from the entire respiratory phase sequence (see “Methods”). Since every second during the task was classified as belonging to a specific respiratory phase (inhalation or exhalation), an increase in the rate of one phase at any given time implies a decrease in the rate of the other phase at that same time point. The results revealed a significant decrease in the inhalation rate compared with the baseline 2 and 3 seconds before MW reports only in the BF condition (2 seconds: *t*(43) = −2.070, *p* = .044, *d* = −.63; 3 seconds: *t*(43) = −2.318, *p* = .025, *d* = −.71; Fig. 4b). Remarkably, as the moment of awareness approached, the participants were more likely to be in the inhalation phase (0 seconds: *t*(43) = 3.134, *p* = .003, *d* = .96; 1 second: *t*(43) = 2.046, *p* = .046, *d* = .62). These patterns were not observed in the SF condition.

**Fig. 4.**
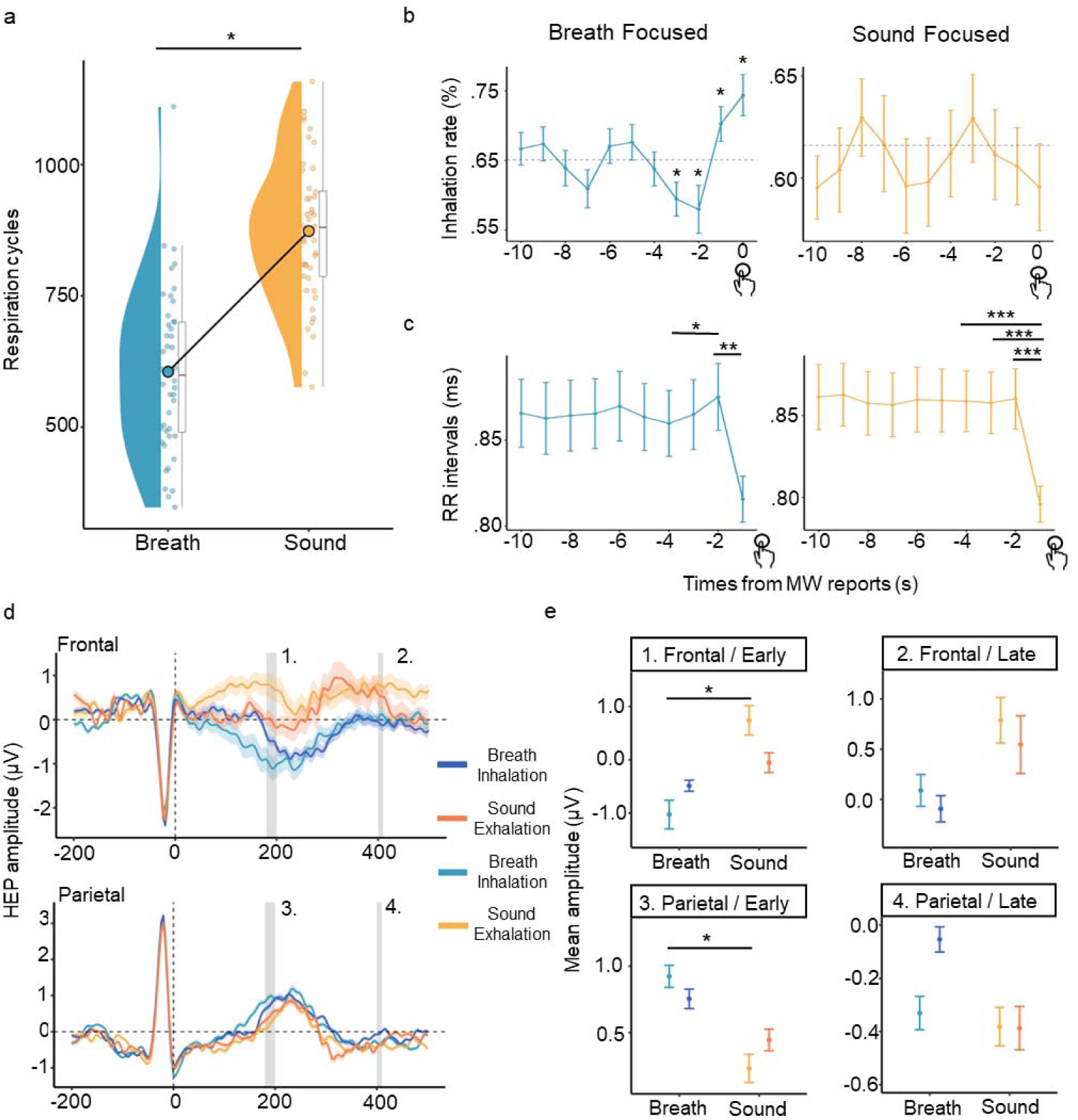
Changes in respiration and the accompanying RR interval and its central process before MW is reported. **a. Number of respiratory cycles during the task:** The task condition is shown on the horizontal axis, and the number of cycles is shown on the vertical axis. Each dot represents a participant. **b. Percentage of the inhalation phase each second before reports:** Inhalation rates are shown on the vertical axis, with the time in seconds (s) from when MW is reported shown on the horizontal axis. The horizontal dotted lines indicate the baseline inhalation rate calculated via random sampling from the entire dataset for each condition. The error bars indicate the standard errors. **c. RR interval at each second before MW reports:** The RR interval is shown on the vertical axis, with the time in seconds from the reports on the horizontal axis. The error bars indicate the standard errors. **d. HEP waveforms during each respiratory phase:** The data from the frontal and parietal electrodes are shown in the upper and lower panels, respectively. The vertical axis shows the amplitude, and the horizontal axis shows the time from the R peak. The bold line shows the mean value, and the range shows the standard errors. The mean HEP amplitude for each participant, focusing on early and late components, was computed by averaging across the periods highlighted in grey. **e. Mean HEP amplitude at frontal and parietal electrodes:** We present the data from the frontal and parietal electrodes in the upper and lower rows. The early component (180~200 ms) and late component (400~410 ms) results are shown on the left and right sides, respectively. The vertical axis shows the mean amplitudes, the horizontal axis shows the task condition, and the colour indicates the respiratory phase. The error bars indicate the standard errors. \**p* < .05, ** *p* < .01, *** *p* < .001.

To explore the effect of the respiratory phases on cardiac activity, we analysed changes in the RR interval over time during periods in which the respiratory phase changed. For this purpose, the RR interval between each second was labelled with the later second—for example, heartbeats that occurred at less than 1 second, between 1–2 seconds, and between 2–3 seconds before MW was reported were labelled 1 second, 2 seconds, and 3 seconds, respectively. To evaluate the RR interval based on the respiratory phase results, the RR intervals were labelled based on time rather than based on the cardiac beat. Through this method, the impact of the respiratory phase at each second is reflected in the RR interval for that second; that is, if multiple heartbeats occurred within 1–2 seconds, both were labelled 1 second. We analysed the RR intervals that occurred within 4 seconds before MW was reported in each condition via a generalized linear mixed model (GLMM), with the timing of the RR interval considered a fixed effect and the participant information considered random effects.

The analysis revealed a significant main effect of the timing of the RR interval (*F*(3, 117.95) = 5.3162, *p* = .002, η*^2^* = .12; Fig. 4c), with subsequent analyses indicating that the RR interval was longer at 2 seconds than at 4 seconds before MW was reported (*t*(118) = 2.817, *p* = .029, *d* = 0.52). The RR interval was shorter at 1 second before MW was reported (*t*(118) = −3.785, *p* = .001, *d* = −.70). This pattern suggests a decrease in heart rate in the exhalation phase, the rate of which increased 3 seconds before MW was reported, followed by an increase in heart rate in the inhalation phase, the rate of which increased 1 second before MW was reported. Similarly, in the SF condition, the main effect of timing was significant (*F*(3, 117.92) = 9.3048, *p* = .002, η*^2^* = .19; Fig. 4c), with comparisons between timings revealing a shorter RR interval 1 second before MW was reported than at 2–4 seconds (2 seconds: *t*(118) = 4.651, *p* <.001, *d* = 0.86; 3 seconds: *t*(118) = 4.254, *p* <.001, *d* = 0.78; 4 seconds: *t*(118) = 4.412, *p* <.001, *d* = 0.81), indicating an increase in heart rate before becoming aware of MW.

In addition, HEP was compared for heartbeats occurring at 0~2 seconds (inhalation) and 2~4 seconds (exhalation) before MW reports to test whether the effects of respiration and heartbeat were reflected in brain activity (Fig. 4d and e). This analysis is preliminary because only half the time window is used compared to the analysis of attentional states, reducing the number of trials. Additionally, we note that the respiratory phase labels were determined based on which phase was increased relative to the baseline in the BF condition. To examine the effects of changes in the respiratory phase on awareness of MW, we labelled each interval as a respiratory phase, although no changes were observed in the SF condition. When only heartbeat data were used for analysis in each phase, the number of trials used to calculate the HEP was further halved. We labelled the data to ensure a larger number of trials and to perform the same analysis as for the RR interval. Although the analysis was performed for a different time window (180–200 ms), we examined the main effect of the respiratory phase and task condition and the interaction effects at the frontal electrode and observed a main effect of the task condition (*F*(1, 216) = 4.700, *p* = .031, η*^2^* = .02). Subsequent analysis revealed an increase in HEP amplitude during exhalation in the BF condition compared with that in the SF condition (*t*(181) = −2.174, *p* = .031, *d* = −. 26). Furthermore, when the same analysis was performed with the parietal electrode data, a similar main effect of task condition was observed (*F*(1, 183.70) = 4.752, *p* = .031, η*^2^* = .03). Subsequent analyses revealed an increase in HEP amplitude during exhalation in the BF condition compared with that in the SF condition (*t*(181) = 2.245, *p* = .026, *d* = .33). No other significant differences were found.

## Discussion

We examined the dynamics of brain activity, the physiological state, and its central process that leads to the transition from MW to the awareness of ongoing thoughts. The results revealed that the HEP amplitude increased when the participants focused on their respiration compared with that in the MW state or when they focused on sound stimuli (Fig. 3c and d). Moreover, the HEP amplitudes were higher when the participants became aware of MW than when they were in the MW state (Fig. 3e and f). No significant changes in RR intervals were observed during the transition from MW to aware states (Fig. 3a), implying that this transition is characterized by the central nervous system processes cardiac activity rather than by physiological states. Additionally, changes in respiratory phases and cardiac activity before becoming aware of MW were observed when the participants were asked to focus on respiration (Fig. 4a). These changes in the respiratory phase before MW reports affect cardiac activity and its central process (Fig. 4c and e).

The HEP is recognized as the central process of cardiac activity^27,28^. Recent studies have revealed that HEP is altered by the attentional resources devoted to processing the internal body state, including respiration^20,34^. Since attention is not necessarily allocated to the body during MW, it is reasonable that the HEP amplitude increased when attention was focused only on respiration (Fig. 3c and d). These results suggest that the increased HEP amplitude before awareness of MW may indicate the increased central process of cardiac activity and the attentional resources directed towards internal states (Fig. 3e and f). Many studies have shown the central process of cardiac activity and conscious experience^7,16,18^. Especially, it is revealed that the self-relatedness of spontaneous thoughts covaried with the heartbeat-evoked responses according to magnetoencephalography (MEG) study^14^. This study revealed that the reactivity of cardiac activity is influenced by the first-person perspective of thoughts. These results suggest that the constant neural update of the visceral states constitutes a self, especially a first-person perspective for conscious experiences^35^. The HEP changes observed in this study may support this hypothesis and reflect that changes in the central process accompany the awareness or changes in conscious experience caused by enhanced central processes of cardiac activity.

We next discuss a potential explanation for the relationship between awareness of MW and the enhanced process of cardiac activity from the perspective of interoception, which is the process by which the nervous system senses and integrates information originating from the body^36^. A previous study on prospective memory—the memory of future intentions—found that cue recognition leads to accelerated cardiac activity for executing intended actions^37^. Furthermore, interoceptive accuracy (IAcc), which is the ability to accurately detect an individual’s internal state, mediates performance in prospective memory tasks^37^. These results suggest that accurate perception of the physiological changes caused by cues enhances processing of the underlying cause and retrieval of the intended action. The self-caught methods inherently involve prospective memory, and the participants need to report, which is required action, whenever they are immersed in MW. As discussed below, the present study and previous studies revealed that various autonomic activities occur with the continuation or leaving of MW^12^. Accurate perception of these fluctuations may promote increased processing related to the underlying event, driving the awareness of ongoing thought. In addition, the dorsal anterior cingulate cortex and anterior insula, which are related to internal and external saliency and interoception^38,39^, showed greater activation just before MW was reported^40^. This result similarly supports a link between the detection of internal changes as salient signals and awareness. From these points, we suggest that not only changes in the central process of the cardiac activity but also the perception of these changes as meaningful contribute to detecting the event, that is, the occurrence of MW.

Based on the respiratory cycle results and the reports of MW in the BF condition, the participants were more likely to be in the exhalation phase before realizing they were in the MW state and in the inhalation phase when they reported MW (Fig. 4b). A previous study revealed that attention to respiration during the exhalation phase led to increased HEP amplitudes^20^. Similarly, it has been shown that the inhalation phase is linked to increased activity in brain areas associated with task engagement, suggesting that inhalation may facilitate increased attention and information processing^41^. These findings suggest that the enhanced central process of cardiac activity in the exhalation phase promotes awareness, leading to reports of MW in the subsequent inhalation phase. Indeed, the heart rate was lower in the interval of increased exhalation than in the interval in which no phase change occurred, and the HEP amplitude was greater during this period than in the SF condition (Fig. 4c and e). These findings suggest that respiration modulation may enhance physiological states and their processing, leading to awareness of MW.

Although we did not observe changes in the respiratory phase in the SF condition, we observed some patterns before MW reports in the BF condition. In the present study, respiration became slower in the BF condition than in the SF condition (Fig. 4a). Furthermore, this slower respiration is associated with changes in cardiac activity and its related processing (Fig. 4c and e), suggesting that attention to respiration enhances the cue signal for awareness of MW. Although further research is needed to fully understand these findings, they provide insights into the mechanisms underlying the benefit of mindfulness. Specifically, meditation practices that focus on breathing are suggested to reduce MW. Meditation extends the duration of the respiratory cycle, leading to deeper and slower breathing patterns^42^. Additionally, as the breathing interval increased, the durations of the longest and shortest heart periods increased and decreased, respectively^43^. These findings imply that mindfulness-based meditation alters the respiratory cycle, enhancing changes in physiological states and its process, which is a cue signal of awareness of MW.

To contextualize these findings within a broader theoretical framework and to explain why these changes in HEP occur, we provide a framework for the transitions in thoughts. Various studies have shown that MW frequency is influenced by motivation for the main task^44–46^. From this perspective, it is suggested that our cognitive resources are limited and that dedicating these resources to one task inherently limits our ability to engage in other tasks, increasing the perceived value of those alternative activities^47^. When an alternative activity is considered more valuable than the current task, boredom may occur, leading to a reallocation of attention towards internal thoughts^48^. This model of MW occurrence could be extended to a background theory of thought shifting; that is, excessive engagement in an MW may lead to a transition to an alternative behaviour other than to internal thoughts due to an increase in aversive emotion for the continuation of MW regardless of the content. A previous study showed that a person being alone with their own thoughts for 15 minutes was so aversive that many participants chose to self-administer an electric shock that they had previously said they would pay to avoid^49^. These results suggest that in situations in which external behavioural options are limited, such as during psychological tasks, participants excessively focus on internal thoughts, leading to aversive emotions and boredom. This relationship was qualitatively reproduced in a simulation, suggesting that excessive engagement in the current state promotes changing action choices^50^. Previous research has suggested that the accumulation of engaging in the same state over time, such as the environment or internal thoughts, may lead to transitions to different thought states.

Next, we discuss the results within the context of our theoretical model. The time series analysis revealed a decrease in alpha wave preceding a reduction in beta wave activity before MW was reported (Fig. 2c). Recent research has suggested that alpha activity is related to internally focused state and MW^32^. We also observed increased alpha activity in the MW state (Fig. 2a). Beta waves are linked to attention control, cognitive load, and stress^51,52^. The decrease in beta activity following the decrease in alpha activity implies that beta activity may be associated with cognitive load and stress levels rather than the regulation of internal thoughts. These results indicate that the decreased alpha activity is associated with leaving the MW state and that decreases in beta activity are associated with decreases in cognitive loads. In an animal study, rats were more actively taking aversive stimuli in an empty environment, with increased activity in the insular cortex immediately before receiving the aversive stimulus^53^. This region is associated with boredom, pain, and interoception, suggesting that the change in action choice occurs when the aversion to the current state and associated physiological response reach their peak. Given that the decrease in alpha activity began 4–5 seconds and ascending neural activity and perceptual sensitivity increased 3.5-5 seconds before MW reports^12,26^, the leaving from MW occurs when the sympathetic activity associated with aversive emotions and related responses peaks. These changes trigger a corresponding increase in perceptual sensitivity and a decrease in the cognitive load. The RR interval increased (2 seconds from reports) after the change in beta activity in the BF condition (Fig. 4c). The negative correlation between the cardiac vagal and beta activity-related brain networks suggests that the release of a high cognitive load leads to increased parasympathetic activity^54^. Multiple subthreshold changes in autonomic activity may occur during the transition from MW to awareness of ongoing thoughts, and these changes may be transmitted to the central nervous system, which is reflected as increases in HEP amplitudes (Fig. 3f). The heart rate tended to increase less than 1 second before MW was reported (Fig. 4c). This result may be regarded as the response to awareness and the readiness state for executing subsequently required actions^37^.

The present study has several limitations. First, while we delineated the relationships between changes in various factors and awareness, the causality of those was not fully understood. For example, we revealed the relationship between changes in alpha and beta activity in terms of temporal continuity. However, it remains uncertain whether leaving the MW state leads to a decrease in cognitive load. To identify the factors that actually drive awareness, such causality must be investigated. In addition, the effects of signals from interoceptive modalities other than cardiac activity were not investigated. Recent research has revealed that the stomach may constrain spontaneous brain activity^55^. Respiration, which was examined only in terms of phase changes in the present study, is also known to influence rhythmic brain activity in multiple frequency bands^56^. Thus, we must consider not only the independent effect of each organ on brain activity but also the interactions between systems. Finally, we could not treat the basic bodily functions in explaining the changes in physiological indices. In the present study, the results of each measure were considered independently in relation to the cognitive process of awareness. However, cardiac activity is known to fluctuate during the respiratory phase, which is known as respiratory sinus arrhythmia^57^. Therefore, changes in cardiac activity may be due not only to cognitive processes, as proposed in the present study, but also to underlying physiological response. Furthermore, these changes may affect central processing via changes in cardiac activity. Although we could not approach to these points because we focused on a short time window of a few seconds and on the gradual cognitive process of awareness, these factors should be viewed within the framework of the autonomic nervous system. In summary, future research should clarify the interactions between the various interoceptive modalities, including their causal effects on awareness, considering influence of basic bodily functions.

In summary, we clarified how brain-body interactions lead to the transition from MW to awareness of ongoing thoughts via EEG and physiological indices. As a result, the leaving from MW occurs due to excessive engagement, and the accompanying changes in physiological states are perceived in the brain, resulting in awareness. This awareness further activates the cardiac activity for subsequent self-reports. Additionally, the modulation of the respiratory cycle via attention to breathing enhanced the cue signals. We suggest that changes in interoceptive processing due to respiratory changes may explain the effects of mindful meditation.

## Methods

### Participants

We recruited 44 healthy university students in Tokyo with normal or corrected-to-normal vision and hearing (24 women; mean age = 21.21; *SD* = 1.96; *range* = 18-29). None of the participants had a history of mental or neurological disorders. This study was approved by the Keio University Research Ethics Committee (No. 18006) and was conducted according to the Declaration of Helsinki. All participants provided written informed consent prior to participation. The participants were paid 7000 JPY.

### Apparatus

In this study, visual stimuli were displayed on a PC monitor placed in front of the participants, whereas auditory stimuli were delivered through speakers located approximately 0.75 metres from either side of the participants at a 45° angle. The participants’ responses were collected via a keyboard.

EEG was recorded during the task using NetStation 5.3.0.1 with a 64-channel HydroCel Geodesic Sensor Net (EGI, Inc. Eugene, OR, USA), referenced to the vertex (Cz). The electrode impedance was maintained below 50 kΩ. ECG data were collected using the NetStation data acquisition system (EGI, Inc. Eugene, OR, USA). In the experiments, Ag/AgCl electrodes were attached to the backs of the right hands and ankles of the participants. The chest wall and abdominal excursions were recorded via inductance plethysmography (Ambu, Ballerup, Denmark and Sleepmate or Perfect Fit 2, Dymedix, St Paul, MN, USA). The presentation of stimuli and the collection of participant responses were controlled via a stimulus delivery and experiment control program (Presentation; Neurobehavioral Systems, Inc., Berkeley, CA, USA).

### Procedure

In the present study, the participants conducted a heartbeat counting task (HCT) and two simple response tasks. We used the HCT to measure participant-specific interoceptive accuracy (IAcc), which is an objective accuracy metric for detecting internal bodily sensations^58^. The analysis of the HCT results is not the main purpose of this study and is therefore included in the Supplementary Materials. Two simple response tasks with the self-caught method were conducted to compare differences in attentional states. EEG, ECG, and respiratory data were measured during these response tasks. Because the response tasks took a long time, they were conducted over two days to reduce participant fatigue. The HCT was always performed on the first day, and the response tasks were counterbalanced among the participants regarding the date on which they were performed first.

### Heartbeat counting task

A pulse oximeter (NONIN, 8600) was placed on the left index finger of the participants. In the HCT, we instructed the participants to silently count their heartbeats at different intervals without taking their pulse orchanging their breathing. In addition, participants were reminded to respond with the number of times they actually felt a heartbeat rather than guessing. This task was repeated six times with time windows of 25, 30, 35, 40, 45, and 50 s, presented in randomized order. The participants began counting after hearing “start” and stopped counting after hearing “stop”. After counting, the participants reported the number of heartbeats they counted and their confidence in their answers via a keyboard. This task was conducted using “cardioception,” a Python package^59^. The absolute proportional difference between the number of reported and actual heartbeats was computed to quantify the cardiac IAcc (Eq. 1).

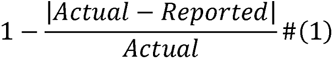

### Two simple response tasks with the self-caught method

The participants performed the two simple response tasks in a dimly lit, soundproof room, and EEG, ECG, and respiratory data were recorded. The task included a sound stimulus response condition (sound-focused; SF) and a breathing response condition (breathing-focused; BF).

In the SF condition, the participants were asked to press the key immediately after each continuously presented pure tone. We randomly changed the interstimulus interval between 1.2 and 2 seconds to prevent habitual responses. Fifty pure tones of various frequencies (range: 400–1400 Hz; interval: 20 Hz) were used to reduce the influence of habituation on the stimulus response. Each sound was presented for 100 ms. The tone was presented at 65 dB. In the BF condition, the participants were asked to press the key after each end of exhalation. The pure tones were also presented in the BF condition for consistency with the environment in the SF condition.

The participants were instructed to report instances of MW whenever they were aware of their thoughts drifting away from the task at hand, guided by the explanations provided before the task^21^. In the present study, MW was defined as the occurrence of thoughts unrelated to the task and stimulus^3,60^. To facilitate their understanding and recognition of MW, we presented examples of both task- or stimulus-related thoughts, such as “How many tones were presented?” and “This tone is C,” as well as thoughts unrelated to the task and stimulus, such as “What shall I have for lunch today?” and “Something upsetting happened yesterday.”

The participants responded to the sound stimuli or the end of exhalation by pressing the “1” key on the keyboard and reported MW with the “3” key. After MW was reported, we asked two questions to capture the content of the participants’ thoughts. First, regarding the emotional valence of the thoughts, the participants used the keyboard to indicate whether their thoughts were negative (“4”), neutral (“5”), or positive (“6”). Second, concerning the temporal aspects of their thoughts, the participants indicated whether these thoughts included the past (“4”), present (“5”), or future (“6”). If the thoughts lacked a temporal aspect or if the participants were uncertain, they were instructed to press “8”. After responding, the participants resumed the task at their own pace by pressing the Enter key. The analysis of the thought content during MW is not the main purpose of this study and is therefore included in the Supplementary Materials.

Each task condition lasted approximately one hour, including rests between the four 12-minute sessions. In addition, a “+” symbol was displayed in the centre of the screen, and the participants were asked to look at the symbol during the task. A schematic representation of the task workflow is presented in Fig. 1a.

### Analysis

#### The definition of attentional states

In our study, we categorized participants’ experiences into three distinct states: MW (MW state), transitioning to awareness (aware state), and focused attention on the main task (focus state). Several previous studies using self-report methods suggested that the participant’s mind wandered even approximately 9 seconds before MW was reported^23–25^. Furthermore, previous studies revealed that the change from MW to awareness begins 3–5 or 4 seconds before MW is reported^12,26^. On the basis of these results, we designated the 5 to 9 seconds before the self-reports as the MW state and the 1 to 5 seconds before the self-reports as the aware state. We excluded the second immediately before the report to reduce the influence of myoelectric potentials arising from MW reports. The focus state was defined as beginning when the first sound stimulus was presented after the task was resumed and last for 4 seconds, as illustrated in Fig. 1b.

To ensure the spontaneity of the MW reports, we placed no limits on the number or timing of MW reports. However, if participants made multiple reports quickly, the same time window was labelled both the MW and task state. We excluded MW reports that occurred within 10 seconds of the preceding report in the EEG analysis. This approach was taken to ensure that the attentional state was accurately identified.

#### Physiological data

##### Respiratory data

All physiological data were preprocessed using MATLAB R2023b (MathWorks Inc. Natick, MA, USA). Respiratory data were used to evaluate associations between the timing of awareness and the respiratory phase. Raw data preprocessing and respiratory phase detection were performed via the BreathMetrics toolbox^61^. Before the raw breathing flow recordings were decomposed into phase data, measurement noise and signal drift were removed. First, the signal was mean-smoothed by a 25-ms window. Next, global linear drift was removed by subtracting the slope of the linear regression model of the data. Local signal drifts were corrected to continuous, minute-long sliding mean baseline windows, and padding was removed. These processes and parameters are defaulted in the package. The respiratory phase was then detected at each time point on the basis of the processed respiratory data.

In this study, respiratory data from the chest and abdomen were acquired to collect data from thoracic and abdominal breathing participants. Since thoracic breathing involves less abdominal movement, we selected the respiratory data that showed more cycles for each participant for subsequent analyses. To investigate the respiratory phase before MW was reported, we employed a one-sample t test. This test was performed to assess whether there was a significant increase in the rate of the inhalation phase before MW was reported compared with that at other time points. Considering that the balance between exhalation and inhalation is unequal under natural conditions, we sampled 10000 data points from the entire respiratory phase. The percentages of inhalation were .651 and .615 in the BF and SF conditions, respectively, and we tested whether the inhalation rate every second before MW was reported was more or less than the baseline value in each condition. The inhalation rate was calculated by dividing the number of inhalations at a given point in time by the number of MW reports. Additionally, we used a paired t test to compare the number of respiratory cycles between task conditions to help elucidate how the task type affects breathing patterns.

##### Electrocardiogram

ECG data were used for HEP calculations and to investigate physiological changes depending on the attentional state. For this purpose, we extracted R peaks from raw ECG signals via HEPLAB^62^. HEPLAB is an EEGLAB extension for automatically detecting cardiac-related events from raw ECG signals. We first used the ecglab slow function included in the package to identify R peaks. Since this function cannot identify all the peaks and may incorrectly identify noise showing abrupt changes such as R peaks, we corrected misidentifications via visual inspection. The R peak timing data were used as events for the HEP calculation.

#### Electroencephalography

##### Preprocessing

All EEG data preprocessing, HEP calculations, and time-frequency analyses were performed via EEGLAB v2023.1 for MATLAB^63^. First, all the acquired raw data were preprocessed for HEP and time-frequency analysis. The recording frequency was 500 Hz, but since the frequency to be analysed was lower than this frequency, the data were downsampled to 250 Hz. Afterwards, continuous EEG signals were filtered via a 0.5–50 Hz bandpass filter. The raw data were processed with an automatic channel rejection method, and power line fluctuations at 50 Hz were removed via the Cleanline EEGLAB plug-in^64^. The data were re-referenced to an average reference computed using the average signal at all EEG electrodes. The EEG signal was then decomposed into 65 independent components via the infomax independent component analysis function included in EEGLAB, and each component was assigned its probability of being the signal source by the IClabel function^65^. We used these labels to remove signals with a probability of 80% or greater of being myoelectric, eye movement, cardiac movement, line noise, or channel noise, which are all recognized as noise. Finally, windows that included noise greater than 30 µV were collected.

##### Heartbeat-evoked potentials

The HEP is an event-related potential based on the R peak that is thought to reflect the central process of cardiac activity. The R peaks were used to segment the preprocessed EEG signals into 800 ms epochs with a −200 ms baseline period. The anterior cingulate cortex, insular cortex, and right frontal electrode have been identified as key areas for HEP source signals^28,66^, with additional evidence suggesting the involvement of the parietal region in processing cardiac activity and spontaneous activity^14^. Accordingly, we focused on electrodes in the right frontal and parietal regions, selecting specific electrodes based on a topographical map (Fig. 3b).

Generally, the early component (200~300 ms) and the late component (400~500 ms) are targeted as the time intervals for evaluating HEP. Although the unified interpretation is still under debate, it has been inferred that the early component reflects perception, whereas the late component is related to higher-order cognitive processing^28^. In the present study, each participant’s average potentials at these electrodes in the 240~260 ms and 400~410 ms intervals from the R peak were used as the HEP data for that individual. The inclusion criteria for the HEP analysis were participants who submitted ten or more reports of MW, ensuring a substantial amount of data to facilitate reliable calculation of the HEP. As a result, six participants who reported fewer than ten instances of MW were excluded from the analysis.

##### Time_frequency analysis

We compared brain activity in different attentional states by examining the spectrum power of different frequency bands throughout the brain. Using a fast Fourier transform (FFT) with a 10% Hanning window, we calculated the spectral power for each EEG channel within this 2–40 Hz range.

We applied a cluster-based permutation test to identify significant activity differences across brain regions and frequency bands in the focus, MW, and aware states^67^ via the FieldTrip toolbox^68^. With this statistical method, the type I error rate caused by multiple comparisons is controlled via nonparametric Monte Carlo randomization. First, cluster-level test statistics are estimated on the basis of the original data and several shuffled versions of the dataset. The cluster-level test statistic is the sum of t values with the same sign across adjacent electrodes and/or frequencies above a specified threshold (i.e., 97.5 quantile of a T distribution). Then, the cluster-level statistics from the original data and the null distribution emerging from the random partitions are compared. The cluster-corrected p value is the proportion of random partitions for which the cluster-level test statistic exceeds that obtained on the basis of the original (non-shuffled) data. The significance level for the cluster permutation test was set to 0.025 (corresponding to a false alarm rate of 0.05 in a two-sided test) ^67^. We conducted a paired-sample t test to compare the following conditions: BF vs. SF conditions in the focus state; focus vs. MW state for each task condition; and MW vs. aware state for each task condition. Our analysis revealed no significant clusters when we assessed the differences between task conditions in the focus and MW states.

All the statistical analyses, excluding the cluster-based permutation tests, were conducted using the open-source software R (R 4.3.1). We used GLMMs to evaluate differences between task conditions and attentional states, as well as to evaluate interactions between states, and were adjusted for the effect of each participant. Simple between-group differences and subsequent analyses for interactions were adjusted by introducing participant information as a random variable. The p values for multiple comparisons were corrected via Tukey’s HSD method. These procedures were performed via the lme4 package in R^69^ GLMM estimation and the emmeans package in R for multiple comparisons^70^.

#### Time series analysis via a statistical model

##### Statistical modeling

We performed a time series analysis to identify the timing and temporal sequence of changes in the spectrum power in each frequency band. For this purpose, the time-frequency analysis method described above was used, and the time window was set to 10 seconds before the MW reports were reanalysed rather than the time windows of attentional states. We then constructed a time series model using a state-space model, integrating change point detection to identify when EEG components shifted from the MW state to the aware state. We converted time units into time steps to perform the analysis. To enhance clarity, we reverted the time steps into actual time units in seconds after this analysis. The window for the time-frequency analysis was slightly shortened from 9.4 seconds before to 0.5 seconds before MW was reported.

The state-space model consists of two separate models: a system model and an observation model. The system model represents the true state, which cannot be measured directly, and the observation model represents the probabilistic distribution of the observed values. In this study, the spectrum power without noise is considered the true state, which is estimated from the observed values. We denote the true state as *µ* and the observed values as *Y*, with *n* indicating participant number and *t* symbolizing the time unit, which was converted from the actual time using the sampling frequency (eq. 2).

Our focus was to identify when the spectrum power began to decrease, indicating a transition between attentional states. We define the *µ* parameter at time *t* as *µ* at the previous time *t*-1 plus a trend component, *δ* (eq. 3). Furthermore, we assumed that this *δ* is the same across individuals and varies randomly over time (eq. 4). For the trend change, we set 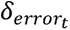, a parameter indicating the strength of the change to *σ_δ_*, which indicates the range of change to implement the time evolution. Assuming that trends in the time series change before and after a change point, we constructed two series of states, *µ_pre_* and *µ_post_*, which started from the same value but evolved differently. That is, before the change point, *Y* is generated according to *µ_pre_*, and after the change point, *Y* is generated according to *µ_post_*. The log likelihood is then estimated for the change points at each time point, and the point with the highest value is estimated as the point at which the trend changes.

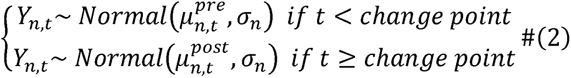

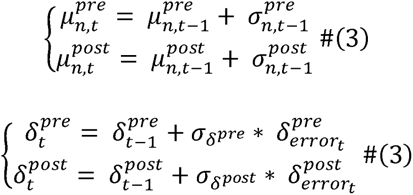

While acknowledging that the mean spectrum power and its noise component can differ among participants, we assume a shared trend of change across individuals. This shared trend underpins our analysis, enabling us to study the overarching pattern of spectrum power changes, irrespective of individual variances. In particular, we considered a half-Cauchy distribution for the observed noise and the distributions of the parameters (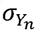 and *σ_δ_*) representing the time evolution of the trend. The half-Cauchy distribution represents the right side of the two symmetric halves of the Cauchy distribution and is recommended as a prior distribution of variance, especially for estimations performed with small samples^71,72^.

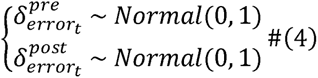

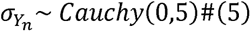

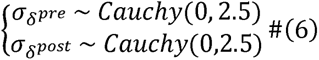

*Model implementation*. The statistical model was estimated via the Bayesian approach using R and the R package Rstan^73^. We employed the Markov chain Monte Carlo (MCMC) method for parameter estimation. In the MCMC method, parameters are estimated by accumulating numerous samples from the posterior distributions. We obtained 6000 simulated samples from the posterior distribution for each parameter. The simulated samples were preceded by 1,000 burn-in samples, which were discarded from the analysis. The burn-in samples were discarded to exclude the effects of the initial values. The MCMC chain was thinned by including only every second draw to reduce temporal autocorrelations among the samples, yielding 2,500 simulated posterior observations. We conducted four samplings and obtained 10,000 simulation data points for each parameter.

The convergence of the MCMC algorithm across the four chains was assessed for each parameter using *R̂* as the index of the variance between the chains relative to the variance within a chain^74^. If the MCMC algorithm converges across all chains, *R̂* is close to 1.0. To estimate the convergence of the algorithm, we set the criterion to be less than 1.1. In summary, across all the parameters, *R̂* values of approximately 1.0 indicated convergence across the four chains. Therefore, the MCMC algorithm converges for all the parameters included in the present model.

## Data and code availability

The data and code that support the findings of this study are available upon request from the corresponding author [K. S.]. The data are not publicly available because we have not obtained consent from the participants.

## Acknowledgements

We are grateful to all the participants in this study.

## Author contributions

All the authors collaborated to develop the study design. K.S. programmed the computer tasks and collected and analysed the data. All the authors interpreted the results. K.S. drafted the manuscript. All other authors provided critical revisions. All authors approved the final version of the manuscript.

---

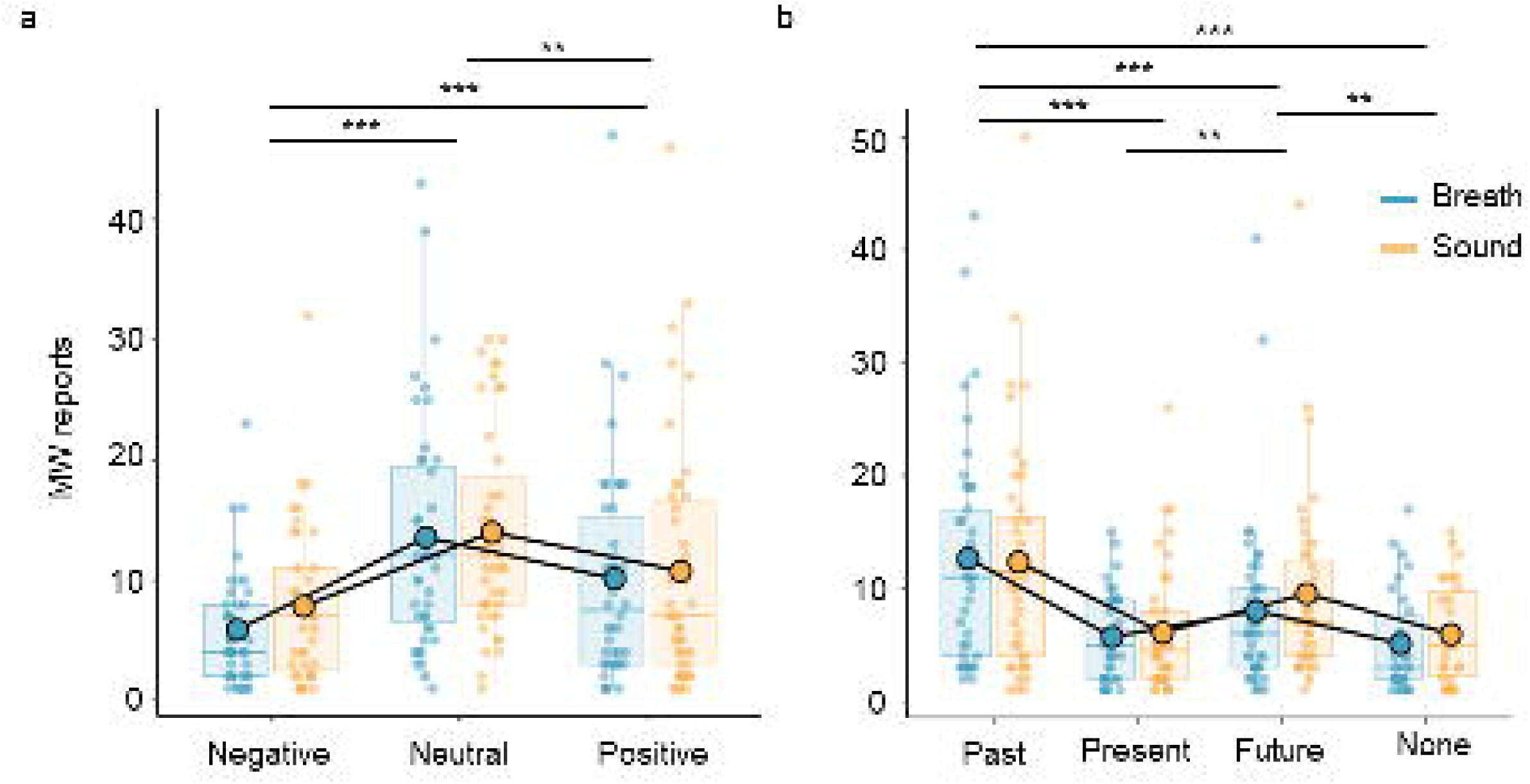

