## Supplementary for "Brain-Body Interactions Influence the Transition from Mind Wandering to Awareness of Ongoing Thought"

### Supplementary Materials

### Results

#### *Mind wandering frequency in each task condition*

We assessed how the frequency of MW reports and the types of thought content varied across task conditions. Although the frequency of thought types during MW has been reported in many studies, the results differ depending on the method used^1-5^. We presented the frequency of thoughts in our experiment to improve our understanding of the nature of MW.

The average number of MW reports was 28.39 (*SD* = 19.22, range = 3-96) in the BF condition and 31.70 (*SD* = 20.51, range = 4-90) in the SF condition (Fig. 2a). The frequency of MW reports tend to be higher in the SF condition than in the BF condition (*t*(43) = -1.880, *p* = .0668, *d* = -.57). The MW frequency is known to vary with task difficulty^6^. The difference in MW frequency in each condition in the present study may be due to the difficulty of responding to an abstract object, such as breathing, to which we usually do not pay much attention.

Next, we examined the thought content during MW and identified a significant main effect of emotional valence (*F*(2, 200.85) = 28.0133, *p* < .001, *η^2^* = .22), with neutral, positive, and negative thoughts occurring more frequently in that order (Supplementary Table 1 and Supplementary Fig. 1a). Additionally, a main effect of time was observed (*F*(3, 260.71) = 26.3036, *p* < .001, *η^2^* = .23), indicating a predominance of thoughts related to the past and future over thoughts related to the present or thoughts without specific temporal features, with past-oriented thoughts being notably frequent (Supplementary Table 1 and Supplementary Fig. 1b).

**Supplementary Table 1. Statistics on differences in the frequency of thought types**

| **Type** | **Between** | **df** | **t value** | **d** | **p value** |
| --- | --- | --- | --- | --- | --- |
| Emotion | Nega-Neutral | 202 | -7.483 | -1.05 | <.001 |
|  | Nega-Posi | 201 | -4.031 | -0.57 | <.001 |
|  | Neutral-Posi | 200 | 3.545 | 0.50 | .0014 |
| Time | Past-Now | 260 | 7.809 | 0.97 | <.001 |
|  | Past-Future | 260 | 4.230 | 0.52 | <.001 |
|  | Past-None | 263 | 7.157 | 0.88 | <.001 |
|  | Now-Future | 261 | -3.678 | -0.46 | .0016 |
|  | Now-None | 264 | 0.024 | 0.003 | 1.000 |
|  | Future-None | 264 | 3.391 | 0.42 | .0044 |

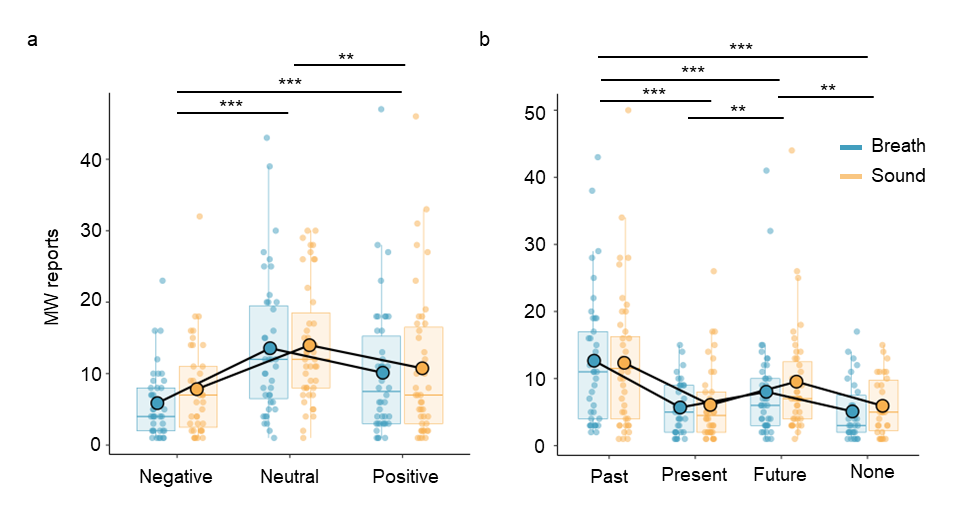

**Supplementary Fig. 1. Frequency of MW reports during the task**

**a, b. Thought content frequency in different task conditions:** The vertical axis shows the frequency of MW reports, and the horizontal axis shows the thought content. Each dot represents the data for one participant. **p* < .05, ** *p* < .01, *** *p* < .001.

***Interoceptive accuracy and mind wandering tendencies***

In our study, in addition to exploring brain-body interactions on awareness, we investigated the relationships between MW and interoceptive accuracy (IAcc), an objective accuracy metric for evaluating the detection of internal bodily sensations^7^. We measured the IAcc via a heartbeat counting task (HCT). A Pearson correlation analysis was conducted to assess the correlation between the frequency of reported thoughts and the IAcc values, adjusting for task conditions (Supplementary Table 2). The findings revealed no significant correlation between the IAcc values and the thought content or MW frequency in this study.

**Supplementary Table 2. Statistics on the correlation between interoceptive accuracy and the frequency of thought types**

| **Type** | **Variable** | **Correlation** | **P value** |
| --- | --- | --- | --- |
| Emotion | Negative | .229 | .139 |
|  | Neutral | .0106 | .946 |
|  | Positive | -.146 | .351 |
| Time | Past | .004 | .981 |
|  | Now | .159 | .309 |
|  | Future | -.073 | .642 |
|  | None | -.059 | .707 |

***Mind wandering contents and physiological indices***

The secondary aim of our study was to investigate the influence of thought content on the physiological state and the HEP. We compared the RR interval for each thought content and task types, but no significant differences were found related to emotion. However, considering the temporal aspect of the thought content, we found the significant main effect of temporal aspcet, task condition, as well as an interaction effect (*F*(3, 254.62) = 3.005, *p* = .030, *η^2^*= .03; *F*(1, 254.63) = 7.903, *p* = .005, *η^2^* = .03; *F*(3, 254.38) = 2.160, *p* = .093, *η^2^*= .02). Subsequent analysis revealed that in the BF condition, heartbeats were slower when the thoughts had no specific temporal aspect than when the thoughts had specific temporal features (*t*(255) = -3.032, *p* = .014, *d* =-.38; t(254) = -3.191, *p* = .008, *d* = -.40, *t*(254) = -3.505, *p* = .003, *d* =-.44). Furthermore, in the absence of a specific temporal aspect, there was a significant difference in the RR interval between the BF and SF conditions (*t*(254) = 3.345, *p* < .001, *d* =.42).

When HEP amplitudes were calculated for each thought content, only the main effect of emotional valence was significant (*F*(2, 73.277) = 3.502, *p* = .035, *η^2^* = .09) within the early component at the frontal electrodes. Subsequent analysis revealed that positive thoughts were associated with reduced HEP amplitudes compared with neutral thoughts, indicating distinct the central process of cardiac activity depending on the emotional valence of thoughts (*t*(72.2) = 2.445, *p* = .044, *d* =.10). No other relationships between HEP amplitude and thought content were found.

### Discussion

In our discussion, we proposed that the physiological changes and the perception of such changes led to awareness of MW. While it might be expected that individuals with higher IAcc scores would be more aware of MW and thus report MW more frequently, our results did not support this hypothesis. This inconsistency may occur due to an unconscious transition from MW to an aware state, which is proposed in our framework. According to our model, over-engagement of the MW state occurs before transitioning to the aware state, a shift that does not rely on signals from the body. This mediation of leaving the MW state may be related to the lack of correlation between IAcc scores and the awareness of MW. These differences also highlight that interoception is especially vital to conscious awareness.

Additionally, the lack of a relationship between MW content and IAcc scores, as derived from behavioral metrics, could be due to the nature of these indicators. The frequency of thoughts as an individual measure might reflect not only awareness tendency but also the inherent frequency of MW, which is unique to each individual. Therefore, confounding may occur, as it is challenging to determine whether an individual inherently experiences less MW or lacks awareness of it. Additionally, while many studies have used the HCT to evaluate IAcc scores, a recent study highlighted the limitations of this task^8^. One of the main problems is its narrow focus on attention to heartbeat, which may not reflect general interoception. Refinement of these measures could offer more precise insights into the link between awareness and interoception.

The decoupling hypothesis suggests that engagement in internal, task-unrelated thoughts reduces the processing of external stimuli during task performance^9^. Studies examining this hypothesis on the basis of the brain's oxygen metabolic energy suggest that the brain's overall energy use is limited, with resources used for MW increasing and decreasing when the perceptual load is low and high, respectively^10^. Despite advancements in delineating the impact of MW on external stimulus processing, how interoception and thoughts interact remains underexplored. The present study revealed that the processing of cardiac activity is reduced during MW in tasks requiring attention to respiration (Fig. 3d), indicating that interoceptive signals may serve as unique perceptual modalities and that the processing of this information may vary during MW. Despite the decreased processing of cardiac activity, the amplitude of these signals fluctuates with thought content. This result implies that the signals from the body are utilized even during MW as a response to emotional aspects that are closely related to interoception^11^.
